# Salient objects in a scene benefit from enhanced perceptual processing and memory encoding: Evidence for a spatial von Restorff effect

**DOI:** 10.1101/2025.03.25.645146

**Authors:** Peiyao Xi, Rongqi Lin, Geoffrey F Woodman, Benchi Wang

**Affiliations:** Center for Studies of Psychological Application, South China Normal University, Guangzhou, China; Institute for Brain Research and Rehabilitation, South China Normal University, Guangzhou, China; Department of Experimental and Applied Psychology, Vrije Universiteit Amsterdam, Amsterdam, the Netherlands; Department of Psychology, Vanderbilt University, Nashville, Tennessee, United States

**Keywords:** visual search, salience, memory, inverted encoding model, phase-amplitude coupling

## Abstract

Rapidly detecting salient objects from surrounding environments is crucial for survival. Our study demonstrates that salient objects in visual search arrays trigger von Restorff-like effects. In a search task, participants detected a tilted target bar among distractors with EEG recordings. The results revealed that salient objects elicited the largest and earliest N2pc component, reflecting early attentional selection, which enhanced multivariate decoding of target location. Importantly, early selection of highly salient targets (25° tilt) triggered a cascade of preferential processing downstream, marked by stronger P3b components, neural synchronization, and phase-amplitude coupling between low- and high-frequency activity, along with better recall performance of target orientation. The strength of memory-related activity on the current trial predicted the vigor of the next selection event, indicating that salience-driven learning influences future attentional control. Overall, object salience in spatial arrays drives a cascade of processing, facilitating rapid learning of object relevance while humans search their environment, similar to the classic demonstrations of the von Restorff effect, except the objects are distributed in space rather than time.

## Introduction

The ability to rapidly detect salient events is essential for survival, allowing us to quickly identify and react to potential threats or opportunities in our environment. This skill may be rooted in our evolutionary history, where recognizing important stimuli—such as predators, food sources, or social signals—was crucial for survival and thriving (Itti & Koch, 2001; Öhman & Mineka, 2001). Neurophysiological research in non-human primates has shown that the brain can respond with remarkable speed and efficiency, exhibiting neural responses to salient signals within 45-107 ms (Buschman & Miller, 2007; Cosman et al., 2018; Katsuki & Constantinidis, 2012; Klink et al., 2023) showing remarkable speed and efficiency of processing. Research in humans has shown that we can use these signals to rapidly learn which locations are likely to contain salient targets versus locations that are to-be-ignored because they are likely to contain salient distractors (B. Wang & Theeuwes, 2018a, 2018b, 2018c; S. Wang et al., 2023; S. Wang & Woodman, 2024).

Here we aimed to test the novel hypothesis that salient objects trigger a von Restorff-like cascade, in which early sensory processing of objects that are salient by being unique in their context is enhanced relative to other stimuli, followed by preferential late-stage electrophysiological activity associated with memory processing of these objects (Fabiani & Donchin, 1995). Such a cascade of preferential processing could explain how humans can rapidly learn to attend to certain salient items, while ignoring others (B. Wang & Theeuwes, 2018a).

The von Restorff effect is the name given to the observation that individuals are more likely to remember items that stand out because of their perceptually unique relative to their context (Von Restorff, 1933), a finding initially described in list-learning tasks (Fabiani et al., 1990; Gratton & Fabiani, 2003; Karis et al., 1984; Rangel-Gomez & Meeter, 2013). In these tasks, participants are given lists to remember and then perform recall. If one of the items on the list is shown in red and the rest in black, memory is better for the perceptually unique items (i.e., the salient objects). This effect has been observed across a wide variety of materials and experimental designs (Wallace, 1965). We hypothesize that the salient item in a spatial array triggers the same kind of cascading von Restorff effect, except playing out across the space of the visual search array instead of the time of the list. In this context, we propose that an object’s uniqueness may arise not only from its temporal isolation (e.g., as in a sequence of items, typical of previous von Restorff experiments) but also from its spatial distinctiveness in a search array. Evidence for such a mechanism could provide a theoretical framework for understanding how humans rapidly learn to attend to or avoid salient items within a cluttered visual environment.

Understanding how salience is handled by the brain is essential as it provides insights into how our brains prioritize and process task-relevant input in real-time, balancing speed and accuracy. To test our von Restorff hypothesis, in Experiment 1, we recorded the electroencephalogram (EEG) from humans performing a visual search task, in which participants were required to search for a tilted bar (with possible target tilts of 3°, 5°, 7°, or 25°) among other vertical bars while keeping fixation at the center of the screen. Behaviorally, they responded to the presence or absence of the target (Figure 1A; see Methods for the details). The 25° target was the most different from its context, producing the most salient signal, whereas the 3° target was less so, as shown in Figure 1.

**Figure 1.**
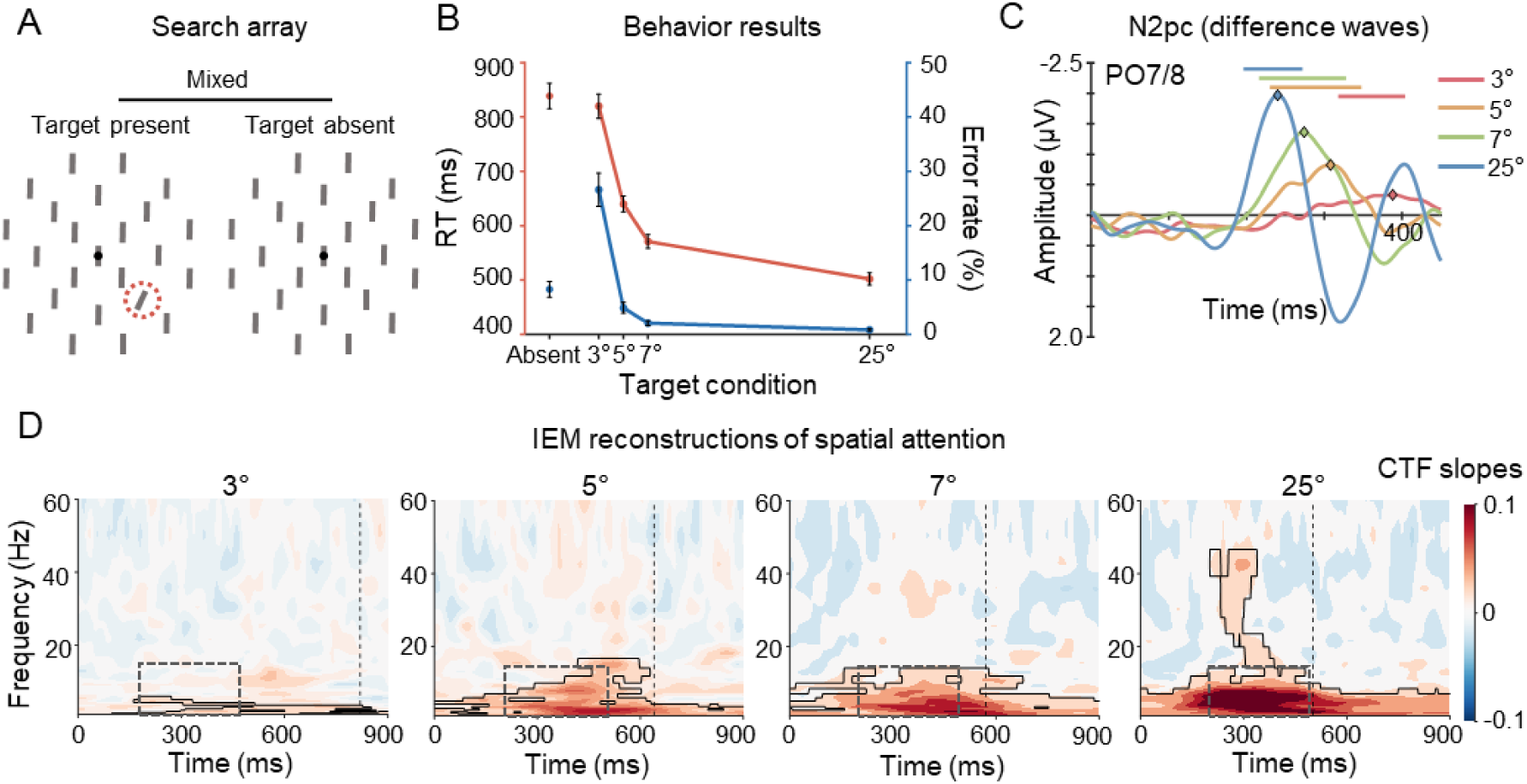
A) Examples of search display. Participants determined whether a tilted bar (3°, 5°, 7°, and 25°) was presented in the search array. The left panel shows a scenario with the 25° target situated in the lower right sector (highlighted with a red dashed circle), while the right panel depicts the target absent condition. B) Behavior results of reaction times (RTs) and error rates. The orange line represents RTs, while the blue line represents error rates. Error bars indicate ±1 standard error of mean (SEM). C) N2pc components were calculated by subtracting ipsilateral activity from contralateral activity toward the target. Significant time windows for N2pc components are highlighted by solid lines (cluster-based permutation test, *p* < .05). Diamond-shaped markers represent N2pc peaks. D) Spatial representations of different targets were tracked using an inverted encoding model (see Methods for details). Significant CTF slopes are covered by solid lines (cluster-based permutation test, *p* < .05). Black vertical dashed lines represent RTs for the respective conditions. Dashed squares highlight the time-frequency window used for comparisons between target conditions.

The von Restorffian hypothesis predicts that we should see effects of salience across stages of processing. However, many theories of visual processing propose that the difficulty of perceptual processing is resolved at that stage of perceptual processing, with the output of perception being essentially invariant to the difficulty of the prior stage (Miller, 1988, Pashler, 1998, Wolfe, 2007). In our case, the subsequent stage of processing that we are going to observe are the potential neural metrics of storage. The alternative hypothesis we will test is the process-purity hypothesis in which perceptual salience is accounted for at the perceptual stage by allowing for more rapid shifts of attention to salient items (Theeuwes, 2018; Treisman, 1985). This predicts that we should observe early perceptual attention effects (i.e., the N2pc) without observing amplitude modulations extending to the metrics of the memory processes that we measure. Instead, the benefits are simply due to attention dwelling slightly longer on such items. If this is true, then we should see N2pc duration should be the best predictor of subsequent memory.

Overall, we predict that the more salient targets (i.e., those with larger angles of deviation) will summon attention effectively, resulting in enhanced memory encoding of these distinct elements, with the learning on the current trial predicting how strongly that salient item will summon attention on the next trial. Experiment 2 tested whether participants exhibit the best behavioral performance when recalling the most salient target during surprise memory test trials after they perform visual search.

## Experiment 1

We recorded the EEG signals when participants performed a visual search task, in which participants were required to search for a tilted bar (with possible target tilts of 3°, 5°, 7°, or 25°) among other vertical bars while keeping fixation at the center of the screen.

## Methods

### Participants

Twenty-four participants (18 female, *M*_age_ = 20.3 years, *SD* = 1.6) were recruited from South China Normal University for monetary compensation. The sample size was predetermined based on previous studies (Liesefeld et al., 2016). Written informed consent was obtained from all participants, and the study was approved by the local ethics committee (2020-3-013). None of the participants had a history of neurological or psychiatric disorders, and all had normal or corrected-to-normal vision.

### Stimuli and apparatus

Stimulus presentation and response collection were controlled by a python program using functions from the PsychoPy package. The search display was shown on a LCD monitor (1920×1080 pixels) at a viewing distance of 74 cm. Participants were instructed to sit as relaxed as possible to minimize muscle activity and reduce electrophysiological noise in electroencephalogram (EEG) signals. The search arrays consisted of 25 gray bars (1.3°×0.25°) on a black background, arranged at the center of the screen and along three imaginary concentric circles around a central bar (radii: 2.1°, 4.2°, and 6.3°). The bars in each circle numbered 4, 8, and 12, respectively. A white fixation dot (radius: 0.2°) was always presented at the center of the screen.

### Procedure and design

The search array remained on the screen for two seconds or until participants made a response. Participants indicated target-present or target-absent responses using the left or right index finger, respectively. If the target was presented, they pressed “x”, if absent, they pressed “m”. If a response was incorrect or no response was made, feedback was shown for 1,000 ms. Trials were separated by an inter-trial interval, randomly jittered between 700 ms and 1,000 ms. Participants were instructed to respond as quickly and accurately as possible. The example of search arrays are showed in Figure 1A.

On one third of trials were target absent trails (all bars vertically oriented), and the remaining two-thirds were target present trials, with the target bar tilted to either the left or right by 3°, 5°, 7°, or 25°. The target was located at one of the eight positions in the second circle of the search array. Different target conditions were mixed. The entire experiment was divided into two sessions, conducted on consecutive days. The first session began with a practice stage consisting of 15 trials. Each session included 16 blocks, each containing 96 trials. Practice trials and trials with incorrect responses were excluded from all the analyses.

### EEG recording and preprocessing

EEG data were recorded using 64 Ag/AgCl active electrodes connected to BrainAmp amplifiers (Brain Products, Munich, Germany), positioned based on the extended 10-20 system, and sampled at 1000 Hz. Specifically, electrodes T8 and FT10 were used to record the vertical electro-oculogram (EOG), electrodes T7 and FT9 were used to record the horizontal EOG, electrodes TP9 and TP10 were used to record signals from mastoids, and electrode FCz was used as on-line reference.

The data were down-sampled to 500 Hz and underwent re-referencing to the mean of the left and right mastoids and were subjected to high-pass filtering with a cutoff frequency of 1.5 Hz (for independent component analysis [ICA] only) and 0.1 Hz (for subsequent analyses). Continuous EEG recordings were epoched from -2000 to 3000 ms relative to the search array onset. Malfunctioning electrodes were visually detected and temporally removed from the data; and a 110-140Hz bandpass filter was used to capture muscle activity and allowed for variable z-score cutoffs per participant based on the within-subject variance of z-scores. Following trial rejection, ICA was conducted solely on the clean electrodes, and components capturing eye blinks, eye movement, or other non-brain-driven artifacts were visually inspected and removed, alongside the vertical and horizontal EOG signals. Subsequently, we interpolated the malfunctioning electrodes identified earlier.

### N2pc, N2, and P3b

Prior to the ERP analysis, preprocessed EEG data underwent bandpass filtering (1-30 Hz) and baseline correction (with a baseline interval of -200 to 0 ms).

N2pc analyses were performed using the PO7 and PO8 electrode sites. The N2pc component was calculated by subtracting the ipsilateral waveform from the contralateral waveform toward the target location to create difference waves over time. The significant N2pc components were determined with cluster-based permutation tests (see below section). For each participant, the N2pc mean amplitude was computed as the average value within the corresponding significant clusters.

N2 and P3b analyses were performed using the Cz electrode site (Rangel-Gomez & Meeter, 2013). After visually inspecting the raw ERP waveforms, we identified the 50% peak latency for both the N2 and P3b components within the time windows of 200-400 ms and 400-600 ms, respectively. To account for potential individual variability, the jackknife procedure (Miller et al., 1998) was applied, systematically excluding one participant at a time and recalculating the 50% peak latency for each iteration. The resulting latencies were then averaged across participants. For the N2 component, the mean amplitude was calculated within a 100 ms window following the 50% peak latency, while for the P3b component, the time window extended 250 ms after the 50% peak latency.

### Time-frequency analysis

To extract power and phase from the time-frequency domains, we selected a set of electrodes for further analysis, including P7, P5, P3, P1, Pz, P2, P4, P6, P8, PO7, PO8, PO3, PO4, POz, O1, O2, CP5, CP1, CP6, CP2, CP3, CPz, and CP4.

Preprocessed EEG epochs were decomposed into time-frequency representations using custom MATLAB scripts. The epochs were convolved with Morlet wavelets, spanning frequencies from 1 to 60 Hz in 25 logarithmically spaced steps. The number of cycles of each wavelet was logarithmically spaced between 3 and 12 to balance temporal and frequency precision. Raw power values were used for the analyses of Inter-trial phase coherence and Phase-amplitude coupling, while they were Z-scored by subtracting the average value and dividing by the standard deviation across all trials for Inverted encoding model analysis.

### Inverted encoding model

To reconstruct spatial representation of target locations, we applied an inverted encoding model (IEM) to estimate spatial channel-tuning functions (CTFs) from the topographic distribution of oscillatory power across electrodes over time (Foster et al., 2017). The assumption is that the oscillatory activity recorded from each electrode reflects the weighted sum of eight spatially selective channels, each tuned to a specific angular position (corresponding to the possible locations of the targets). The response profile of each spatial channel across target locations was modeled as a half sinusoid raised to the seventh power:

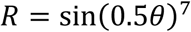

where θ is the angular location (ranging from 0° to 359°), and R is the response of the spatial channel in arbitrary units. To ensure that the peak response of each spatial channel was centered over one of the eight target locations, the response profile was circularly shifted.

We partitioned the data randomly into independent sets of training (2/3trials) and test data (1/3 trials) following a cross-validation routine. The training data (B; m electrodes × n locations) were used to estimate the weights that approximated the possible contributions of the eight spatial channels to the observed oscillatory activity measured at each electrode and target location. We defined C (k channels × n locations) as a matrix of the predicted response of each spatial channel (estimated by the basic function for that channel) for each location; and W (m electrodes × k channels) as a weight matrix characterizing a linear mapping from “channel space” to “electrode space”. We further described the relationships between B, C, and W in a linear model as follows:

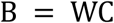

Using the weight matrix (W) obtained via least-squares estimation, we inverted the model to transform the observed test data into estimated channel responses (i.e., reconstructed CTFs). This IEM routine was iterated 10 times to minimize the influence of idiosyncrasies specific to any trial assignment and obtain averaged channel-response profiles. We then calculated the slope of the CTFs via linear regression to quantify attentional processing for different target locations over time. Higher slope values indicate greater spatial selectivity while lower values indicate less spatial selectivity. Target-tuning CTFs were reconstructed according to the target location across participants and was applied to each time-frequency point.

### Inter-trial phase coherence

Inter-trial phase coherence (ITPC), which measures the degree of consistency in instantaneous oscillatory phase across trials at different frequencies and time points, was examined separately for the 25° and 3° target. Additionally, the differences between the 25° and 3° target were also calculated. ITPC estimates were computed from the complex time-frequency representations obtained via wavelet convolution, as described in the Time-frequency analysis section. ITPC across trials was calculated using the following formula:

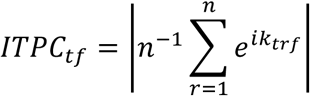

where *n* is the number of trials and *k* is the phase angle at time-frequency point *tf* for trial *r* (Cohen, 2014). ITPC is the magnitude of the mean vector formed by all phase vectors, each with a length of 1, across all trials at a given time-frequency point. ITPC values range from 0 to 1, where 0 indicates no phase consistency across trials, and 1 indicates perfect phase consistency across all trials.

### Phase-amplitude coupling

The phase-amplitude coupling (PAC) between low-frequency (1–15 Hz) phase and high-frequency (20–60 Hz) power for each electrode was examined using mean vector length (Canolty et al., 2006). It was calculated according to the following equation:

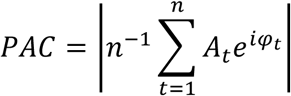

where *n* is the number of data points, *A_t_* is the power at data point *t*, and *φ_t_* is the phase angle at data point *t*. The raw PAC values were then assessed against random distributions via permutation testing. For this, we selected the 200-500 ms time window for PAC analysis. Specifically, the time series of high-frequency power was shuffled, and this process was repeated 200 times to create random distributions of PAC values. The raw PAC values were then z-scored based on the random distributions to correct for any spurious results. The z-scored positive PAC values were compared to 0 using paired t-tests, with significant results considered where the PAC values were significantly larger than 0.

### Cluster-based permutation test

To correct multiple comparisons, we used a cluster-based permutation test against a null-distribution shuffled from 1000 iterations (following Monte Carlo randomization procedure). Specifically, t-tests were performed across participants for conditional differences against zero to identify above-chance activity, and time windows with *t*-values larger than a threshold (*p* = .05) were combined into contiguous clusters based on adjacency. The cluster statistics was defined as the sum of the *t* values within each cluster. The null distribution was formed by randomly permuting condition labels for 1000 times to get the largest clusters per iteration. Clusters were determined to be significant if the cluster statistics was larger than the 95th percentile of the null distribution.

### Correlation analysis

We calculated the Spearman correlation coefficient between the coupling strength on the current trial and the N2pc peak of the next trial. Specifically, we focused on clusters where the PAC difference between the 25° and 3° target was significant, particularly in the 4-7 Hz range modulating 33-60 Hz, as depicted on the rightest side of Figure 3A. For each trial, we computed the raw coupling strength within this frequency range for both the 25° and 3° target conditions, excluding any coupling strengthen values that were more than 2.5 standard deviations away from the mean. Subsequently, we identified the N2pc peaks for each trial within the corresponding significant N2pc clusters for both target conditions, taking into account that the mean amplitude varied considerably across trials.

## Results

Behaviorally, mean response times (RTs) and error rates as a function of the salience of the target are presented in Figure 1B. A repeated measures ANOVA with target condition (target absent, target tilts of 3°, 5°, 7° or 25°) as a factor revealed significant main effects for both mean RTs, *F*(4, 23) = 186.15, *p* < .001, *η_p_^2^* = 0.89, and error rates, *F*(4, 23) = 69.9, *p* < .001, *η_p_^2^* = 0.75. Further comparisons showed that mean RTs for the 25° target were significantly faster than for other target conditions, as well as the target absent condition, all *p*s < .001. Additionally, mean error rates for the 25° target were significantly lower compared to the other conditions, including target absent, all *p*s < .001. Detailed follow-up comparisons are provided in Tables A1 and A2 in the appendix. These results indicate that the 25° target generated more salient signals, leading to faster responses and fewer errors. In contrast, the 3° target was less salient, resulting in slower responses and more errors.

### Early salience-driven selection: N2pc component

Our first measure was the N2pc component of the participants’ EEG waveforms, as this component is believed to index the earliest shift of attention toward targets in search arrays (Eimer, 1996; Woodman & Luck, 2003). To determine whether target salience changed how perceptual attention was focused, we analyzed this lateralized event-related potential (ERP; see Methods for details). A strict cluster-based permutation test was employed to identify the precise time window for the N2pc component across conditions, allowing us to measure expected latency shifts of the N2pc across target salience which should be observed if attention shifts more slowly to some targets than others. As shown in Fig 1C, for the 25° target trials, a reliable N2pc was observed between 196-274 ms (cluster-based permutation test, *p* < .05), indicating remarkably fast responses to salient signals. With each step down in target salience, the onset of the N2pc component was delayed. Specifically, we observed a significant N2pc for the 7° target from the interval of 214-330 ms, for the 5° target from the interval of 228-350 ms, and for the 3° target from the interval of 312-414 ms. Additionally, we observed differences across target conditions for mean amplitude within corresponding clusters, *F*(3, 23) = 37.86, *p* < .001, *η_p_^2^* = 0.62. Further comparisons showed that mean amplitude for the 25° target was significantly larger than for other target conditions, all *p*s < .001. Detailed follow-up comparisons are provided in Tables A3.

In summary, we observed the earliest neural responses toward the most salient targets (i.e., the 25° target), starting at about 196 ms. These results support the conclusion that salient targets benefit from preferential early perceptual processing, but can the brain activity trigged by salience be used to find the target in space? To address this, we applied an inverted encoding model (IEM; see Methods for details) to track target processing over time and space so that we could examine frequency-specific responses across different target conditions.

### IEM reconstructions across target conditions

The IEM model not only detects early attentional allocation but can quantify how useful that information is in telling the organism where the salient object is in the environment. We applied the IEM model over time across a broad range of frequencies (1-60 Hz) using parietal and occipital electrodes. As shown in Figure 1D, regardless of target salience, reliable target selectivity (reflected by channel tuning function [CTF] slopes) was detected across 1-15 Hz and sustained for approximately 900ms after target onset (cluster-based permutation test, *p* < .05). To compare target selectivity following initial detection between conditions, we calculated mean CTF slopes in 1-15 Hz range over 200-500 ms. A repeated-measures ANOVA with target condition as a factor revealed a significant main effect, *F*(3, 23) = 85.85, *p* < .001, *η_p_^2^* = 0.8. Further comparisons showed that mean CTF slopes was significantly larger for the 25° target compared to other conditions, all *p*s < .01. Detailed follow-up comparisons are provided in Table A4. Additionally, only significant CTF slopes for the 25° target were observed specifically in the high-frequency range (15-47 Hz) over 202-388 ms. Together, these findings suggest that the brain continues to process salient signals from the most salient targets (i.e., 25°), with more salience providing clearer information about where the target was in space.

### Memory-related ERPs following the early attentional selection

Classic electrophysiological studies of the von Restorff effect show that salient items in a word list trigger enhanced N2 components, similar to the N2pc effect observed in the current study, and also evoke a subsequent P3 effect (Fabiani et al., 1990; Kamp et al., 2013; Karis et al., 1984). Specifically, unique items in a word list generate a larger posterior P3b (Kamp et al., 2013), with this component related to working memory updating (Donchin & Coles, 1988), and other memory processes (Rugg & Curran, 2007).

Figure 2A shows the N2 and P3b components time-locked to the onset of the search array, along with the differences in mean amplitude across conditions (see Methods for details). These plots show that the most salient items elicited the largest amplitude N2 components, followed by largest amplitude P3b components. Specifically, a repeated-measures ANOVA with target condition as a factor revealed a significant main effect for mean amplitude of N2, *F*(3, 23) = 27.48, *p* < .001, *η_p_^2^* = 0.54; and P3b, *F*(3, 23) = 25.38, *p* < .001, *η_p_^2^* = 0.53. Further comparisons showed that: 1) mean amplitude of N2 for the 25° target was significantly larger than other conditions, all *p*s < .001; 2) amplitude of P3b for the 25° target was significantly larger than 5° target and 3° target, both *p*s < .01. Detailed follow-up comparisons are provided in Tables A5 and A6.

**Figure 2.**
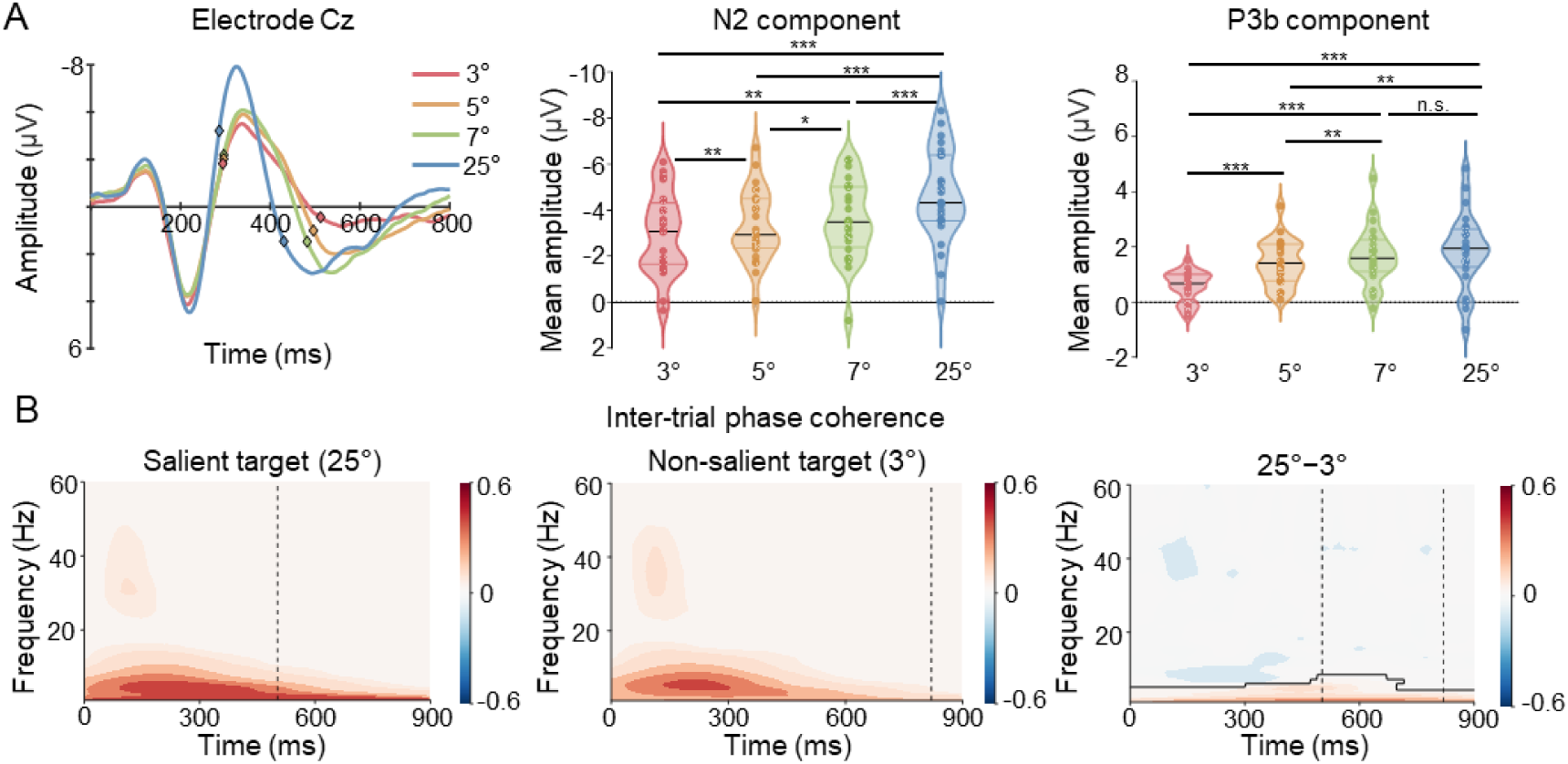
A) The left figure shows ERP components at the Cz electrode, with diamond markers indicating the onset of the N2 and P3b components. The middle and right figures represent the mean amplitude of the N2 and P3b components, respectively. \**p* < .05, \*\**p* < .01, \*\*\**p* < .001; n.s. denotes no significant difference between conditions. B) Inter-trial phase coherence (ITPC), reflecting neural synchronization, for the 25° and 3° target, along with their difference. Significant ITPC differences between conditions are covered by solid lines (cluster-based permutation test, *p* < .05). Black vertical dashed lines represent RTs for the respective conditions.

These findings are strikingly similar to what is observed with unique items in a word list, with such isolates also eliciting enhanced N2/P3b complexes (Kamp et al., 2013; Karis et al., 1984; Rangel-Gomez & Meeter, 2013). This alignment supports the prediction that more salient items elicit stronger downstream memory-related activity as measured by the P3b of subjects’ ERP waveforms. Next, we examined whether these memory effects would extend to measures of neural synchronization.

### Memory-related synchronization following the early attentional selection

If salient items trigger a von Restorff cascade that enhances perceptual process and memory storage of items that are salient due to uniqueness, then we should also be able to see that the downstream memory enhancement on neural synchronization. Foundational theorists proposed that stronger synchronization in the initial period following presentation should result in better long-term memory encoding (Hebb, 2005). If this neural process occurs following enhanced selection of the more salient items, then we should see more coherent activity during the processing of the salient target (25°) compared to the non-salient target (3°). To investigate this, we calculated inter-trial phase coherence (ITPC) for the 25° and 3° target separately. ITPC measures the consistency of neural oscillatory phase across trials at specific times and frequencies, with high ITPC values reflecting well-aligned phases and the engagement of a specific neural network. Conversely, low ITPC values may indicate greater variability in neural responses, potentially due to differences in attentional allocation. If enhanced attention is followed by better memory storage of a salient stimulus, then we expect stronger phase consistency for more salient targets. Our results revealed larger ITPC values when processing more salient targets, indicating stronger phase consistency and more coherent neural synchronization. As shown in Figure 2B, reliable phase consistency (reflected by ITPC values) was observed for both the 25° and 3° targets across 1-60 Hz over the entire selected time window (cluster-based permutation test, *p* < .05). Notably, phase consistency was significantly stronger for the 25° target compared to the 3° target across 1-7 Hz throughout the entire time window. This confirms that the brain effectively and consistently coordinates its activity to process the salient targets, setting the stage for enhanced downstream memory representations of these targets.

### Phase-amplitude coupling after initial target detection

As previously mentioned, we successfully tracked target selectivity for the salient target (25°) across high frequency bands (15-47 Hz) based on frequency power, and observed differences in phase synchronization within low-frequency bands (1-7 Hz) between the 25° and 3° targets. This suggests a potential link between low-frequency phase and high-frequency neural activity during salience processing. To investigate this, we analyzed phase-amplitude coupling (PAC) between the phase of 1-15 Hz oscillations and the power of 20-60 Hz activity within the 200-500 ms window after target onset. As shown in Figure 3A, for the 25° target, we detected reliable coupling strength where power in the 20-47 Hz and 55-60 Hz ranges was modulated by the phase of the 4-7 Hz range, and power in the 47-55 Hz range was modulated by the phase of the 1 Hz and 7-14 Hz range (cluster-based permutation test, *p* < .05). For the 3° target, we observed that the phase in the 4-6 Hz range modulated the power of 23-47 Hz range, the phase in the 4-7 Hz range modulated the power of 55-60 Hz range, and the phase in the 1-1.5 Hz and 7-14 Hz range modulated the power of 47-55 Hz range. Notably, as shown in Figure 2B, the coupling strength elicited by the salient target (25°) was stronger than the non-salient target (3°) within the 33-60 Hz power range, which was modulated by the phase of the theta band (4-7 Hz; cluster-based permutation test, *p* < .05). These results confirm that the brain processes the salient target (25°) by enhancing coupling strength between the theta band and high-frequency activity, likely driven by its salience. This kind of coupling of high and low frequency activity is particularly useful when needed to encode events across multiple neural structures (Fries, 2005), as occurs during long-term memory encoding (Friese et al., 2013).

**Figure 3.**
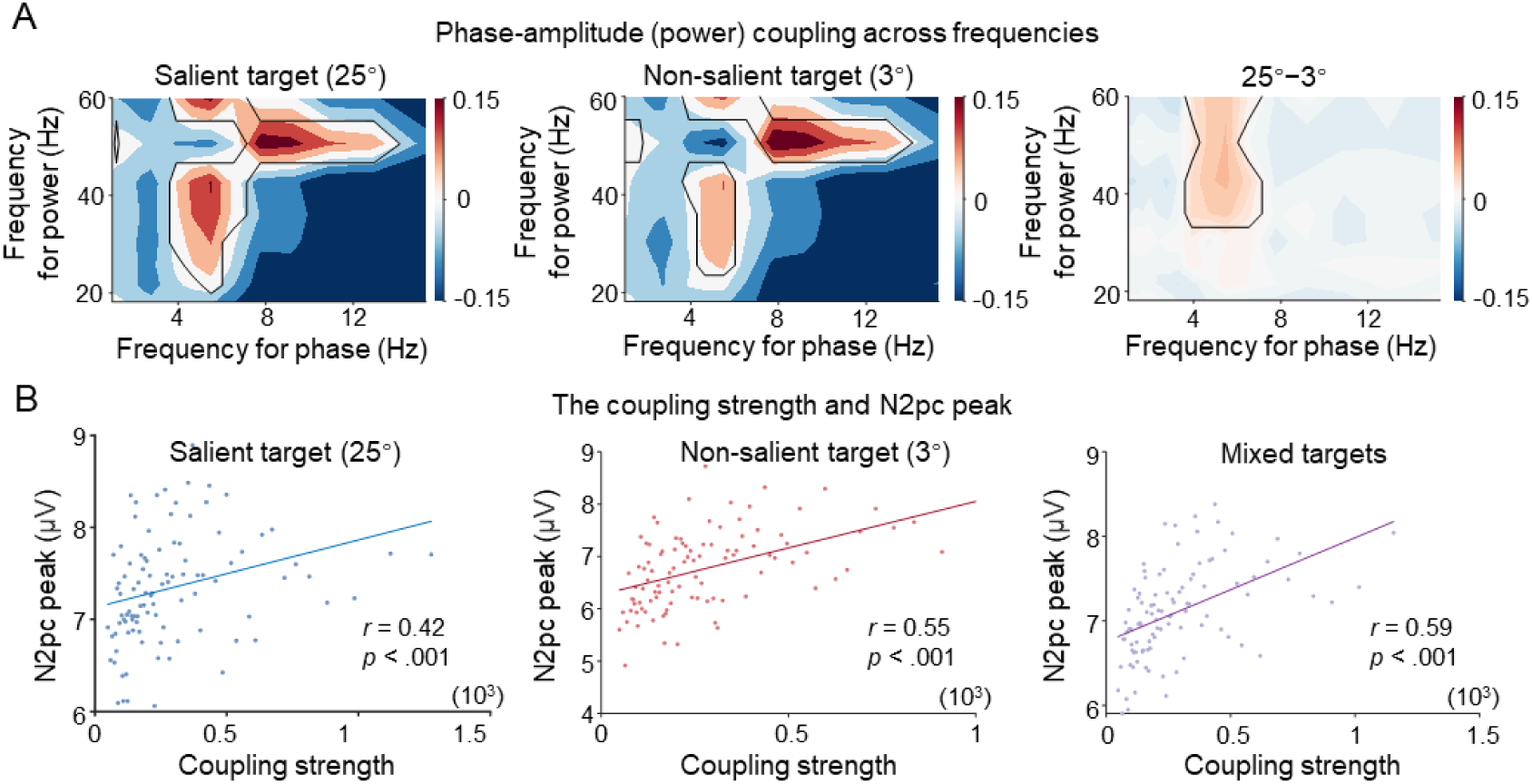
A) Phase-amplitude coupling (PAC) for the 25° and 3° target, along with their difference. Significant PAC values for each condition and significant differences between conditions are covered by solid lines (cluster-based permutation test, *p* < .05). B) The correlation between coupling strength and N2pc peak for the 25° and 3° target, and their mixture.

### Memory-related PAC predicts subsequent attentional selection

Next, we assessed whether the observed memory effects improved processing on subsequent trials. We hypothesized that a von Restorff cascade underlies the rapid learning of salient item predictability during visual search, influencing attentional control on subsequent trials. To test this, we performed a sequential-trial analysis, measuring coupling strength on each trial to determine if it predicted the N2pc amplitude elicited by the same item upon its next appearance. We computed the Spearman correlation between coupling strength on the current trial and the N2pc peak value on the subsequent trial. To minimize trial-by-trial variance, trials were sorted by coupling strength into 100 bins, with data averaged within each bin, yielding 100 observations. The analysis revealed significant positive correlations between coupling strength on the current trial and N2pc peak on the next trial, regardless target type or their mixture (25° target: *r* = 0.42, 3° target: *r* = 0.55, mixed targets: *r* = 0.59, all *p*s < .001; see Figure 3B). Notably, these significant correlations remain robust across various bin selection (e.g., 200, 500, and 1000), and even when using the raw trial data.

Although these correlations are modest, their significance survives the fact that the target salience was randomized, which allowed for many intervening trials between the learning trial and the target’s reappearance. This time course supports our hypothesis that salient items trigger a von Restorff cascade, enhancing memory, which is subsequently used to improve processing on future trials. Specifically, the strength of memory activity on the current trial predicted how strongly attention was biased toward that same item when it reappeared later in the experiment.

## Experiment 2

Our results revealed that early selection of highly salient targets (25° tilt) triggered a cascade of preferential processing, characterized by stronger P3b components, enhanced neural synchronization, and robust phase-amplitude coupling between low- and high-frequency activity. However, one might question whether these neural signatures truly reflect memory and learning, as their behavioral relevance was not directly measured. We addressed this concern in Experiment 2 by having participants perform a shorter version of the visual search task. Then, at the end of the task, they were unexpectedly asked to recall the orientation of the targets, allowing us to assess whether recall performance was better for more salient targets (e.g., 25°). We selected targets with tilts of 3°, 7°, and 25° and assumed that the 7° target, rather than the 3° target, would provide a more meaningful comparison with the 25° target. The rationale was that the 3° target, being very close to the vertical line, might allow participants to complete the task by simply clicking near the vertical position, resulting in the smallest recall errors overall.

## Methods

We recruited 45 participants (30 female, *M*_age_ = 21.9 years, *SD* = 3.8) via the online platform NAODAO (https://www.naodao.com). After providing informed consent, participants completed a visual search task identical to Experiment 1, consisting of 120 trials (two-thirds with a target). Following the search task, participants were unexpectedly asked to recall the target’s direction five times by selecting one of 360 positions on a wheel using a mouse, with no time limit (see Figure 4A). Participants were unaware of the recall requirement during the search task. They were randomly assigned to three groups, searching for the targets with tilts of 3°, 7°, or 25°, respectively. Stimulus presentation and response collection were programmed in PsychoPy 2021.2.3 Builder and exported as a JavaScript file for online testing.

**Figure 4.**
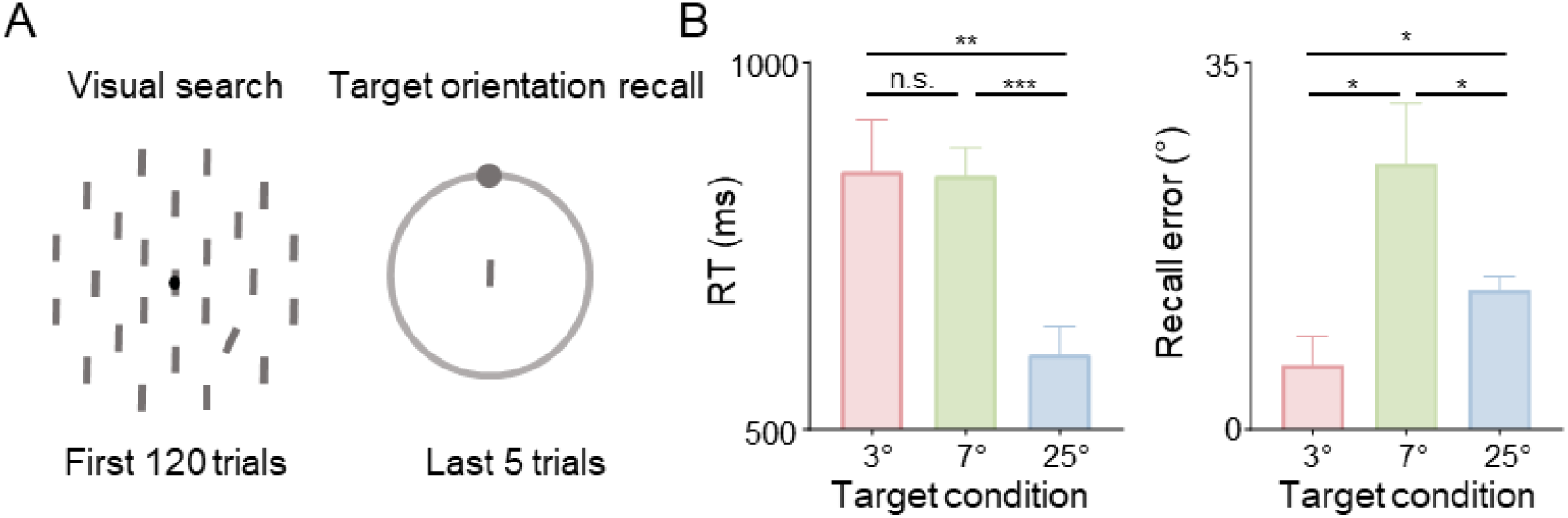
A) Examples of the search and recall displays in Experiment 2. Participants completed a visual search task identical to Experiment 1, comprising 120 trials (two-thirds containing a target). After completing the search task, participants were unexpectedly instructed to recall the target’s orientation five times by selecting one of 360 positions on a wheel using a mouse, with no time limit. Importantly, participants were unaware of the recall requirement during the search task. B) Mean RTs and recall errors across different target conditions. Error bars represent ±1 SEM. \**p* < .05, \*\**p* < .01, \**p* < .001; n. s. indicates non-significant differences.

## Results

Behaviorally, mean RTs and recall error magnitudes as a function of target salience are presented in Figure 4B. These findings are consistent with the prediction that people remember the more salient targets better than the less salient ones when tested with a surprise memory test. We ran separate repeated measures ANOVAs with target condition (target tilts of 3°, 7°, or 25°) on mean RTs and error magnitudes.

These revealed significant main effects for both mean RTs, *F*(2, 42) = 7.71, *p* < .001, *η_p_ ^2^* = 0.27, and recall errors, *F*(2, 42) = 6.69, *p* < .001, *η_p_ ^2^* = 0.24. Further comparisons showed that the mean RTs for the 25° target (601 ms) were significantly faster than mean RTs for the other target conditions (3° target: 851 ms; 7° target: 846 ms), both *t*s > 2.99, *p*s < .009, *d*s > 1.13. No significant difference was observed between the 3° and 7° targets, *t* < 1. Recall accuracy showed a similar pattern to the RT data. Mean magnitude of recall error for the 25° target (13.3°) was significantly smaller than that of the 7° target (25.4°), *t*(28) = 2.15, *p* = .049, *d* = 0.75, indicating that participants remembered the salient target more accurately than the less salient target. Mean recall errors for the 3° target (6.1°) was significantly lower compared to the 7° and 25° target, both *t*s > 2.66, *p*s < .019, *d*s > 0.86. This suggests that participants recalled the 3° target by simply clicking near the vertical position, resulting in the smallest recall errors overall.

## General discussion

The brain must continuously adapt its processing to optimize the detection of behaviorally relevant signals (Bassett et al., 2011). Here we tested the novel hypothesis that current object salience has such strong effects on future deployments of attention due to triggering a von Restorff effect in space, rather than across time as it has been observed in the classic memory literature. Our findings supported the predictions of this hypothesis by showing that the most salient target (25°) resulted in the fastest responses, the lowest error rates, the strongest N2pc components, and the clearest spatial decoding of the target location using EEG patterns, supporting the idea that the brain is especially attuned to salient stimuli. In contrast, less salient targets (3°, 5°, 7°) produced slower responses and weaker neural signatures, emphasizing the differential neural investment required for processing stimuli of varying salience.

If visual salience were a computation that was resolved at early perceptual stages, as is sometimes assumed (Reynolds & Desimone, 2003; Fecteau & Munoz, 2006), then we should have seen our effects were restricted to N2pc amplitude and duration effects, with subsequent memory processes differing only in the time each neural signal was focused on the different items. If item uniqueness in space triggers a processing cascade similar to how the brain handles uniqueness across time (i.e., within word lists), then unique items in spatial scenes would also trigger a von Restorff effect, whereby these items are better stored in memory compared to less salient items. To test this, we focused on the memory-related activity that followed the preferential selection of the salient items. First, we found that the most salient items elicited the largest amplitude P3b components, a neural signature associated with memory processing (Polich, 2012). Second, we found that more salient items also elicited stronger synchronous activity across EEG electrodes, further supporting the idea that salience enhances neural coordination and memory encoding. Next, we detected the strongest target selectivity (as reflected by CTF slopes) across the 1-15 Hz frequency range within 200-500 ms post-target onset, consistent with previous studies linking low-frequency oscillations to memory processes (Riddle et al., 2020; Rodriguez-Larios & Alaerts, 2019). The enhanced target selectivity in the delta band (1-3 Hz) further supports this, as delta activity is associated with memory processes in previous work (Kim et al., 2019).

In our task, highly salient targets (25°) were detected more easily, as indicated by shorter RTs and more precise memories of those objects, making the search process easier for these targets. A reader might be concerned that the observed memory effects are driven by search difficulty rather than salience manipulation. However, it is important to note that easier searches require less processing time by definition, so these salient items are viewed for less time and that would be expected to result in worse memory for those objects. However, we observed the opposite pattern in which the objects that were on the screen for the least amount of time are remember the best.

Moreover, we observed stronger P3b components, enhanced neural synchronization, and robust phase-amplitude coupling between low- and high-frequency activity, accompanied by better recall performance of target direction for the more salient targets.

Importantly, we found that the strength of the learning, measured on the current trial using PAC, was correlated with the amplitude of the N2pc elicited by that same target on the next trial. This conditionalized analysis provides evidence that attentional and memory processes triggered by salience are interconnected, such that learning on the current trial affords enhanced processing on the next trial. Notably, furthermore, the process-purity hypothesis proposes that perceptual salience is accounted for at the perceptual stage by allowing for more rapid shifts of attention to salient items (Theeuwes, 2018; Treisman, 1985). This predicts that we should observe early perceptual attention effects (i.e., the N2pc) without observing amplitude modulations extending to the metrics of the memory processes that we measure.

However, this is not the case in the current study. We observed both N2pc duration and amplitude was associated with salience manipulation. That is, the benefits are not simply due to attention dwelling slightly longer on salient items. Together, these findings suggest that after enhanced perception of salient items, the brain preferentially stores information about those items in memory, thereby optimizing future target selection as we search our environment.

In conclusion, our results suggest that detecting and processing salient targets involves more than a simple detection mechanism. The brain preferentially deploys attention to unique, salient objects, and then stores that information in memory to facilitate improved performance on subsequent trials. Yet what is the nature of the memory mechanisms at play here? It would be tempting to conclude that our measures of synchrony and coupling reflect encoding and storage within working memory, which would be sufficient to support task completion. Notably, this type of temporary memory storage does not appear to account for the full scope of saliency effects during visual search. Specifically, it appears that information about salient objects in search arrays may not even be stored in explicit memory at all. Instead, it may be stored in implicit memory. The idea that implicit memory is driving learning during search tasks like ours comes from evidence that statistical learning in the absence of awareness is ubiquitous (Turk-Browne et al., 2005). The proposal being that the brain incidentally processes all of the properties of salient objects, creating memory traces of learned regularities. These regularities, in turn, can influence behavior even in the absence of explicit memory. Thus, our findings provide support for the idea that salience alters how objects are handled by perceptual and memory systems. Future research will need to determine the nature of the memories that we then use to control perceptual analysis.

## Acknowledgements

PX and BW designed the experiment, PX collected the data, PX, RL and BW analyzed the data, GW and BW wrote the paper. All authors approved the final version of the manuscript for submission. The work was funded by the National Science and Technology Innovation 2030 Major Program (2022ZD0204802), the MOE Project of Key Research Institute of Humanities and Social Sciences in Universities (22JJD190006) and the Natural Science Foundation of Guangdong grant (2023A1515012789) to BW; and the National Science Foundation (BCS-2147064), and the National Institutes of Health to GFW (P30-EY08126) to GW. The codes and behavioral data can be accessed through https://github.com/xipeiyaoo/detect-salient-signals. The EEG data can be made available upon reasonable request.

## Appendix tables

**Table A1.**
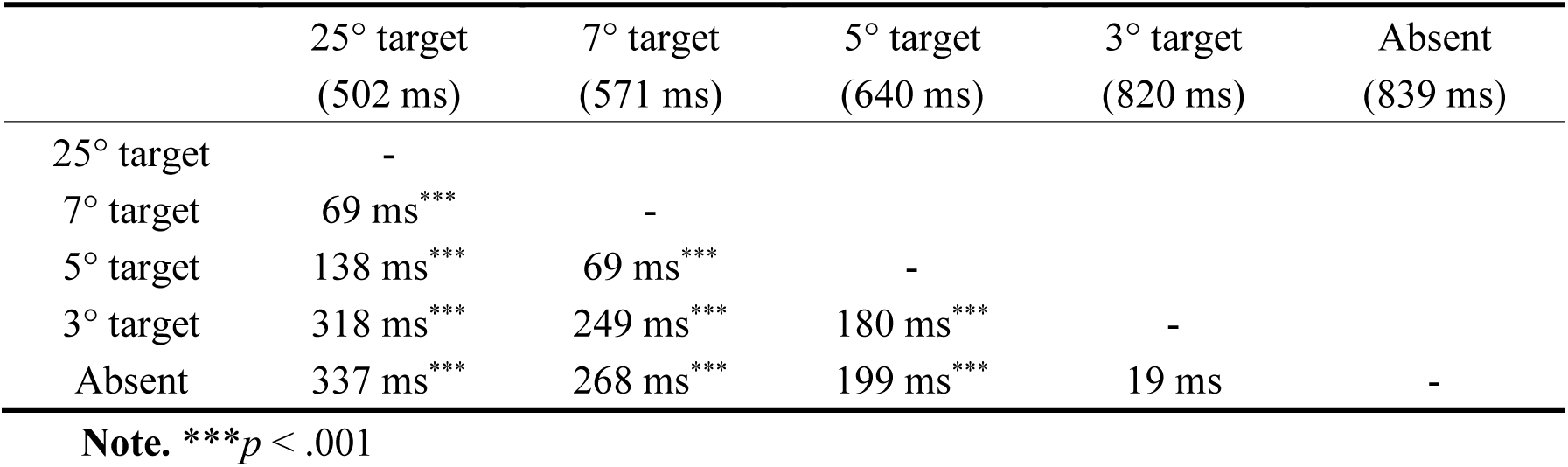
Paired comparisons between conditions (mean RTs)

**Table A2.**
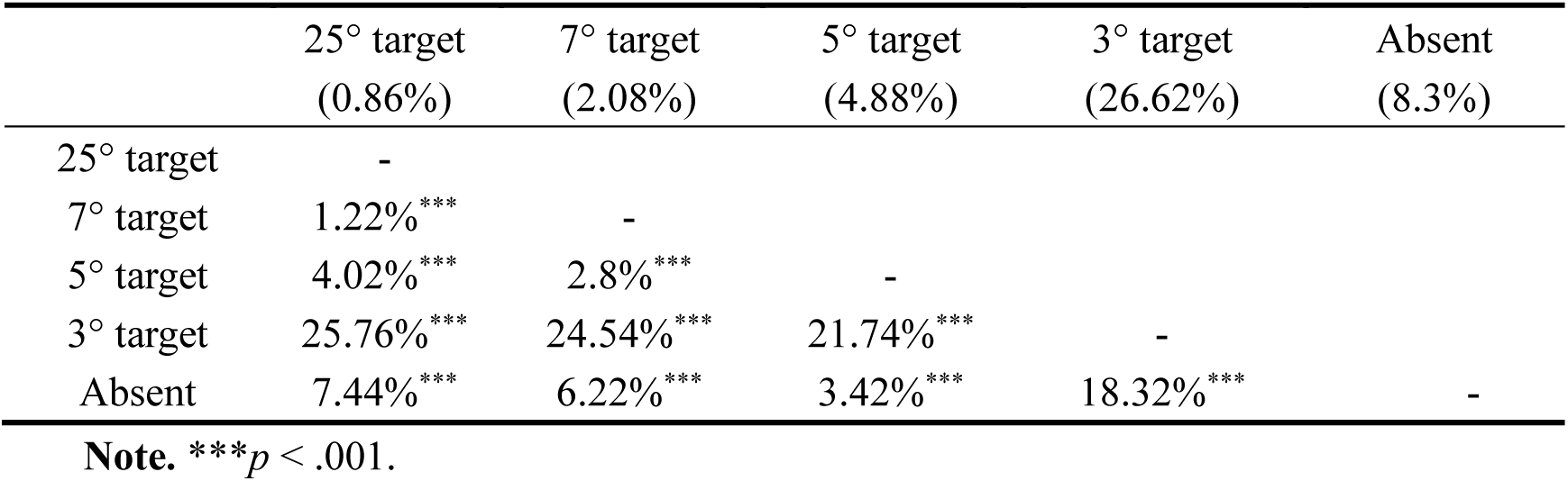
Paired comparisons between conditions (mean error rates)

**Table A3.**
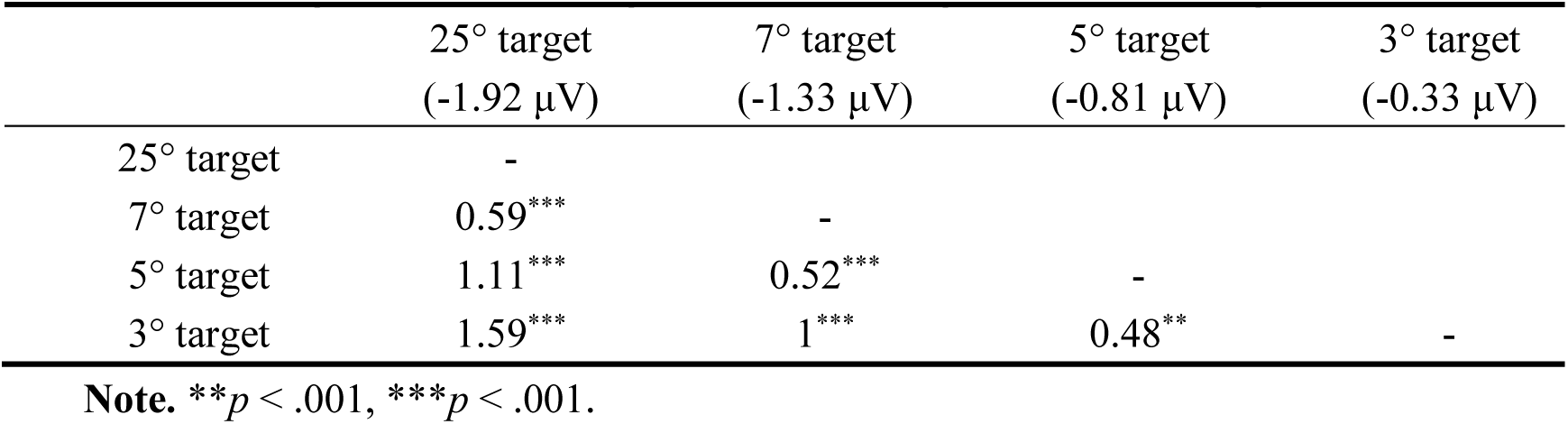
Paired comparisons between conditions (N2pc mean amplitude)

| | 25° target<br>(-1.92 μV) | 7° target<br>(-1.33 μV) | 5° target<br>(-0.81 μV) | 3° target<br>(-0.33 μV) |
| --- | --- | --- | --- | --- |
| 25° target | - |  |  |  |
| 7° target | 0.59<sup>***</sup> | - |  |  |
| 5° target | 1.11<sup>***</sup> | 0.52<sup>***</sup> | - |  |
| 3° target | 1.59<sup>***</sup> | 1<sup>***</sup> | 0.48<sup>**</sup> | - |

**Note.** <sup>\*\*</sup> $p < .001$ , <sup>\*\*\*</sup> $p < .001$ .**Table A4.** Paired comparisons between conditions (average CTF slopes in 1-15 Hz range over 200-500ms)

**Table A4.** Paired comparisons between conditions (average CTF slopes in 1-15 Hz range over 200-500ms)

|  | 25° target<br>(0.081) | 7° target<br>(0.057) | 5° target<br>(0.044) | 3° target<br>(0.018) |
| --- | --- | --- | --- | --- |
| 25° target | - |  |  |  |
| 7° target | 0.024<sup>***</sup> | - |  |  |
| 5° target | 0.037<sup>***</sup> | 0.013<sup>**</sup> | - |  |
| 3° target | 0.063<sup>***</sup> | 0.039<sup>***</sup> | 0.026<sup>***</sup> | - |

**Note.** <sup>\*\*\*</sup> $p < .001$ .

**Table A5.** Paired comparisons between conditions (N2 mean amplitude)

| | 25° target<br>(-4.6 μV) | 7° target<br>(-3.58 μV) | 5° target<br>(-3.33 μV) | 3° target<br>(-2.95 μV) |
| --- | --- | --- | --- | --- |
| 25° target | - |  |  |  |
| 7° target | 1.02*** | - |  |  |
| 5° target | 1.27*** | 0.25* | - |  |
| 3° target | 1.65*** | 0.63** | 0.38** | - |

**Note.** \* $p < .05$ , \*\* $p < .01$ , \*\*\* $p < .001$ .

**Table A6.** Paired comparisons between conditions (P3b mean amplitude)

| | 25° target<br>(1.88 μV) | 7° target<br>(1.67 μV) | 5° target<br>(1.42 μV) | 3° target<br>(0.58 μV) |
| --- | --- | --- | --- | --- |
| 25° target | - |  |  |  |
| 7° target | 0.21 | - |  |  |
| 5° target | 0.46** | 0.25** | - |  |
| 3° target | 1.3*** | 1.1*** | 0.84*** | - |

**Note.** \*\* $p < .01$ , \*\*\* $p < .001$ .

## Notes

**Conflict of interest statement** The authors declare no competing interests.

https://github.com/xipeiyaoo/detect-salient-signals

### Competing Interest Statement

The authors have declared no competing interest.

### Summary of Updates

Updated Title: The manuscript title has been changed from "Salient objects in a scene trigger enhanced perceptual selection and memory encoding" to "Salient objects in a scene benefit from enhanced perceptual processing and memory encoding: Evidence for a spatial von Restorff effect" to better align with the study's core findings and focus. Addition of sample size justification； References updated.

